# R/UAStools::plotshpcreate: Create Multi-Polygon Shapefiles for Extraction of Research Plot Scale Agriculture Remote Sensing Data

**DOI:** 10.1101/2020.02.21.960203

**Authors:** Steven L. Anderson, Seth C. Murray

## Abstract

Agricultural researchers are embracing remote sensing tools to phenotype and monitor agriculture crops. Specifically, large quantities of data are now being collected on small plot research studies using Unoccupied Aerial Systems (UAS, aka drones), ground systems, or other technologies but, data processing and analysis lags behind. One major contributor to current data processing bottlenecks has been the lack of publicly available software tools tailored towards remote sensing of small plots, and usability for researchers inexperienced in remote sensing. To address these needs we created plot shapefile maker (R/UAS::plotshpcreate), an open source R function which rapidly creates ESRI polygon shapefiles to the desired dimensions of individual agriculture research plots areas of interest and associates plot specific information. Plotshpcreate was developed to utilize inputs containing experimental design, field orientation, and plot dimensions for easily creating a multi-polygon shapefile of an entire small plot experiment. Output shapefiles are based on the user inputs geolocation of the front left corner of the research field ensuring accurate overlay of polygons often without manual user adjustment. The output shapefile is useful in GIS software to extract plot level data tracing back to the unique IDs of the experimental plots. Plotshpcreate is available on GitHub (https://github.com/andersst91/UAStools).

## 1. Introduction

Remote sensing platforms geared towards automated high-throughput crop monitoring have become important tools with potential to drive gains in crop improvement and management (Araus et al., 2018). Although curating sensor information/images has become somewhat trivial, especially for remote sensing specialists, processing sensor information into informative data for decision making remains a tedious, time consuming, and challenging process (Shakoor et al., 2017; Shakoor et al., 2019). Aside from the processing/calibration of sensor datasets, reducing dataset dimensionality is a critical step in facilitating the ability to make actionable decisions. In plot-based agriculture research programs this requires the creation of individual areas of interest (AOI) for each research entry/treatment of interest. These AOIs are used to extract plot level information, such as the plant height, canopy cover, or vegetation index of a specific plot containing an individual genotype or experimental treatment. When the number of plots is small (<50), little effort is required and shapefiles containing AOIs can be manually drawn. However, for large plant breeding or genetics programs hundred to multiple thousands of plot AOI may be needed and unique identifying information with consistent and repeatable labeling is needed for each AOI.

There are several features that are needed to make plot extraction from GIS software efficient, even for novices. (i)The ability to rapidly create a grid of polygons to be overlayed on plots in the proper rotation for any mosaic. (ii) The ability to easily incorporate the experimental design using tabular information with attributes, such as, unique plot IDs for each polygon. (iii) An option for buffering, which is important when plots are not immediate adjacent, with some border such as a walkway/alley. Buffering also is useful to reduce plot overlap when an orthomosaic has some distortion. (iv) Free and open source availability that allows all researchers to use the same tool without proprietary software.

Tools available to rapidly create AOI polygons for large scale small plot trials (> 100 of plots) are limited (Table 1), or unknown to the user community. ArcGIS (ESRI Development Team, 2019) and QGIS (QGIS Development Team, 2019) utilize a fishnet approach to create a regular gridded rectangle, although unique identifiers must manually be assigned to each polygon. Unique ID assignment is further complicated due to the left-to-right, top-to-bottom grid creation rather than the bottom-to-top, serpentine design commonly used in small plot design. ArcGIS and QGIS require identification of four-point coordinate system to properly orient gridded polygons to the field-plot offset from north-south orientation. Progeny (Progeny Development Team, 2019), a commercial software, resolves this issue through manual identification of corner plots within your field of interest and saves plot information as range, row information but, lacks a plot identifier. There is still a need for an open source resource that incorporates (i) plot orientation, (ii) experimental design, (iii) automated attribute table with unique plot ID, and (iv) plot buffering.

**Table 1.** Available software which can create gridded multi-polygon shapefiles.

| Software | Function Name | Connects unique ID | Utilized experimental design | Automates buffer zone | Automated Rotation | Open Source |
| --- | --- | --- | --- | --- | --- | --- |
| ArcGIS [4] | Create Fishnet | F | F | F | T | F |
| QGIS [5] | Vector Grid | F | F | F | T | T |
| Progeny [6] | Grid | F | F | T | T | F |
| R/UAStools [7,8] | Plotshpcreate | T | T | T | T | T |

## 2. Implementation

R/UAStools::plotshpcreate is implemented as a software package function of R (Figure 1a), which constructs a multi-polygon shapefile (.shp) of a research trial, with individual polygons defining specific research field plots. Plotshpcreate has two dependency packages (R/sp (Pebesma and Bivand, 2005; Bivand et al., 2013) and R/rgdal (Bivand et al., 2019)) and is recommended to be executed on the most current version of R. Plotshpcreate has three main argument inputs (i) seed preparation and experimental design data frame (Figure 1b), (ii) A-B line coordinates (Figure 1c and d), and (iii) plot and buffer dimensions (Figure 1e). Output files include a multi-polygon ESRI shapefile using overall plot dimension and a multi-polygon ESRI shapefile using buffer plot dimension. Optional outputs include visual representations of shapefile for rapid accuracy assessment. UAStools can be loaded into the R environment using the devtools package (Figure 1a).

**Figure 1.**
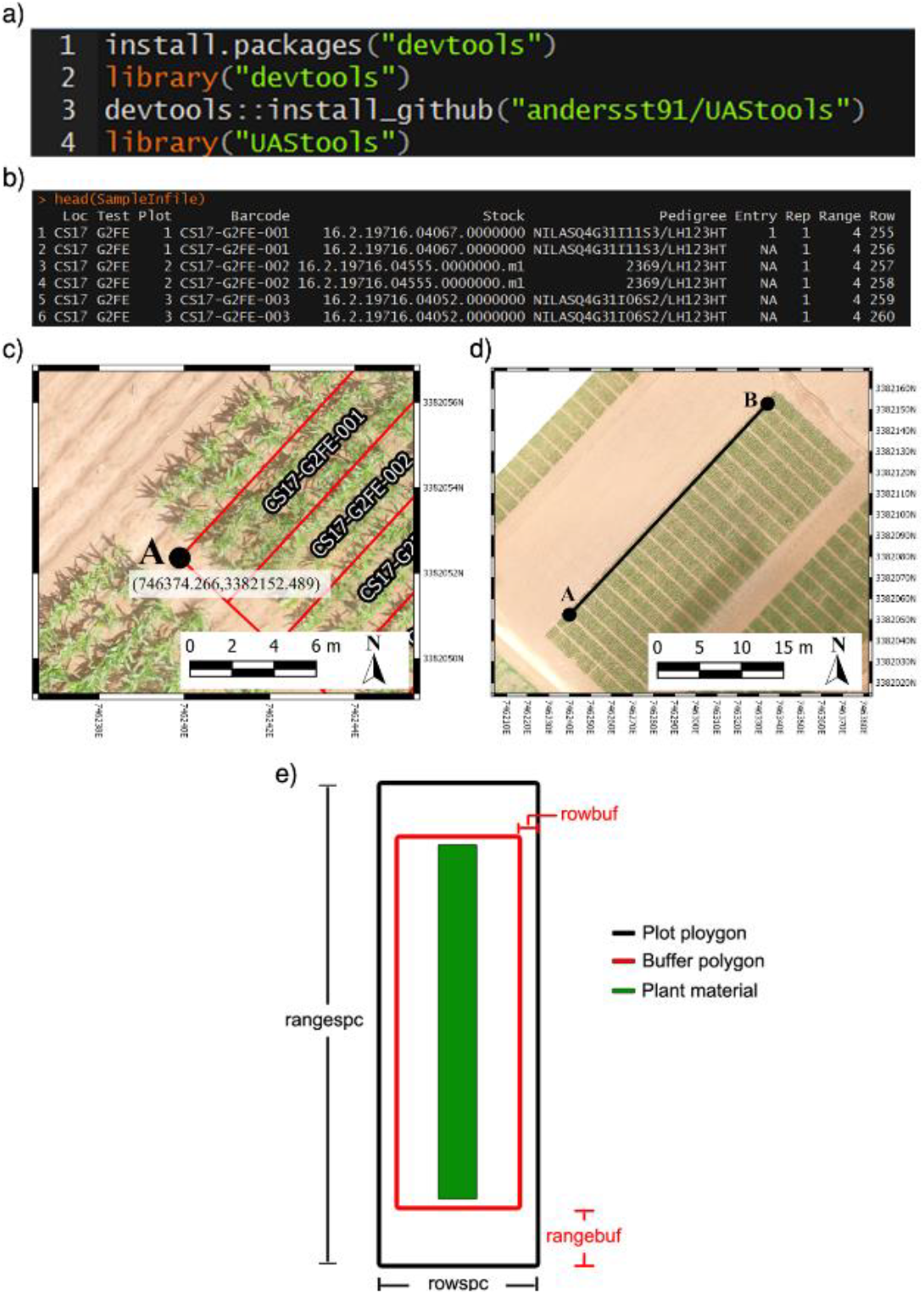
**[a]** Executable lines necessary to load UAStools into the R environment. **[b]** Example of common data structure used as the input file for plotshpcreate. **[c]** Demonstrates the localization of a “A” point in reference to the first plot polygon. **[d]** Visual representation of A-B line. **[e]** Diagram demonstrating the plot (black) and buffer (red) polygon spacing input parameter.

### 2.1 Required inputs

#### 2.1.1. Seed preparation and experimental design data frame

The infile (Figure 1b) for plotshpcreate.R requires four columns matching the quoted column names below (additional columns are permitted but won’t be utilized): (i) “Plot”: The number of each plot (numeric); (ii) “Barcode”: A unique identifier for each plot (character); (iii) “Range”: The range (also called row) number of each plot in the experimental grid (numeric); and (iv) “Row”: The row (also called column) number of each plot in the experimental grid (numeric). Barcodes must be unique across all observations if nrowplot=1 (i.e. if every observation of the infile has a unique barcode use nrowplot==1). Repeated barcodes and plot numbers if there are multi-row plots as the plotshpcreate function accounts for this redundancy within the function. Barcodes must be identical across adjacent rows in a plot if trial consists of multi-row plots. An example from the barcode system we typically use is “CS17-G2FE-018” where “CS” denotes the location, “17” denotes the year, “G2FE” denotes a trial in this location year and “018” denotes the 018^th^ plot within this trial. A sample dataset has been provided with R/UAStools and is defined in R as “SampleInfile” when UAStools is loaded via library("UAStools").

#### 2.1.2. A-B line coordinates

Plotshpcreate was developed for Universal Transverse Mercator (UTM) coordinates. Please convert to UTM before attempting to use plotshpcreate. Many such converters are available online. Plotshpcreate builds plot polygons based on the “A” point (Figure 1c) as a reference and utilized the plot locations in the rectangular grid (Range, Row) of the experimental design to calculate the appropriate geo-locations for the polygon corners. The location of "A" is specific, and must lie at the bottom left corner of the first plot. More specifically, within the middle of the preceding alley and in the middle of the inter row space to the left of the first plot (Figure 1b). The B point is less specific but should be place in the same inter-row space to accurately capture the exact angle (i.e. deviation from South/North orientation) of the field (Figure 1c). The best method for the A-B line development is using the geo-rectified orthomosaic of interest, alternatively a high-confidence a handheld real-time kinematic (RTK) GPS on a pole to ensure an accurate A-B line in the field. If many temporal orthomosaics will be used throughout the season, one of these with high accuracy and low distortion can be used to develop plotshpcreate and subsequently applied to all other timepoints.

#### 2.1.3 Plot and buffer dimensions

There are four polygon dimensions arguments that can be specified to accurately create the proper plot dimensions and buffer dimensions desired (Figure 1e). Row (i.e. column) spacing (rowspc) spacing of a single row is set to 0.76 m in reference to the row spacing, by default. Range (i.e. row) spacing (rangespc) refers to the total plot length including half alley distance on either side of the plot (default: 7.62 m). Row buffer spacing (rowbuf) is the distance removed from both sides of rowspc to create a buffer zone between plots boundaries (default is 0.03 m). Range buffer spacing (rangebuf) is the distance removed from both sides of rangespc to create a buffer zone between plots boundaries (default is 0.61 m). As an example, if alleys are 1.22 m rangebuf should be set to 0.61 m to remove 0.61 m from both ends of the polygon. These settings all must be changed for each researchers plots sizes, any default will almost never fit any other research study.

### 2.2 Optional functionality arguments

We have designed plotshpcreate to have several useful functionalities that dictate how plot polygons can be created (Table 2). Plotshpcreate was developed based on a common style of seed preparation input files, meaning that if a plot consists of multiple planted rows, the input file must contain each row of data for each grid of the planting design (e.g. every range x row combination) with the same unique ID. There are ways to overcome this by adjusting plot dimensions and input file, but we will not discuss those methods.

**Table 2.** Gallery of plotshpcreate input parameters.

| Parameter | Default | Options | Description |
| --- | --- | --- | --- |
| A | NULL | User | Numeric vector of UTM coordinates (Easting,Northing) of "A" point. |
| B | NULL | User | Numeric vector of UTM coordinates (Easting,Northing) of "B" point. |
| infile | NULL | User | Data frame containing seed preparation file and experimental design |
| outfile | NULL | User | Character assignment to define output file names. |
| nrowplot | 1 | >0 | Number of adjacent rows that constitute a plot/unique ID |
| multirowind | FALSE | Logical | Logic parameter that indicates if adjacent plot rows should be combined and treated as a single plot shapefile and unique identifier. |
| rowspc | 2.5 | >0 | Row (i.e. column) spacing of a single row. |
| rowbuf | 0.1 | $\geq 0$ | Distance removed from both sides of rowspc to create a buffer zone between plot boundaries. |
| rangespc | 25 | >0 | Range (i.e. row) spacing of a single row. |
| rangebuf | 2 | $\geq 0$ | Distance removed from both sides of rangespc to create a buffer zone between plots boundaries. |
| stagger | NULL | User | Numeric vector of length three defining [1] row where staggers starts, [2] rows sowed by planted in a single pass, and [3] stagger offset distance from A point. |
| plotsubset | 0 | $\geq 0$ | Defines how many adjacent rows should be excluded from either side of the plot. |
| field | NULL | User | Character vector to indicate the trial the shapefile is being developed for. Example: CS17-G2FE |
| unit | feet | feet and meter | Character vector that the unit of measure for the polygon dimensions. |
| SquarePlot | TRUE | Logical | Logic parameter to indicate if PDF file is desired for visualization of none rotated polygons. |
| RotatePlot | TRUE | Logical | Logic parameter to indicate if PDF file is desired for visualization of rotated polygons. |

The default arguments assume single row / range plots (nrowplot=1) and a unique barcode for each row of the input file (equivalent to Figure 2a). It is common to have multiple adjacent rows plots where researchers desire a single measurement representing the combined rows. Plotshpcreate combines multi-row plots (Figure 2b) based on matching barcodes by defining the number of rows a plot contains (nrowplot=”n”) and telling plotshpcreate to combine the rows (multirowind=F). Plotshpcreate can create single polygons of each row plot of a multi-row plot (Figure 2a), adding an index to each Unique ID in order to identify the data of the multirow plot from left to right (e.g. left row: CS17-G2F-018_1, right row: CS17-G2F-018_2, etc.). Individual row polygons of a multi-row plot can be created with the arguments multirowind=T and defining the number of rows a plot contains (nrowplot=”n”). Multirow plots with rows extracted individually in this way can be averaged after extraction or during analysis. However, while a two row plot (for example) will double the number of observations, these will not be independent and caution should be used in interpretation of degrees of freedom.

**Figure 2.**
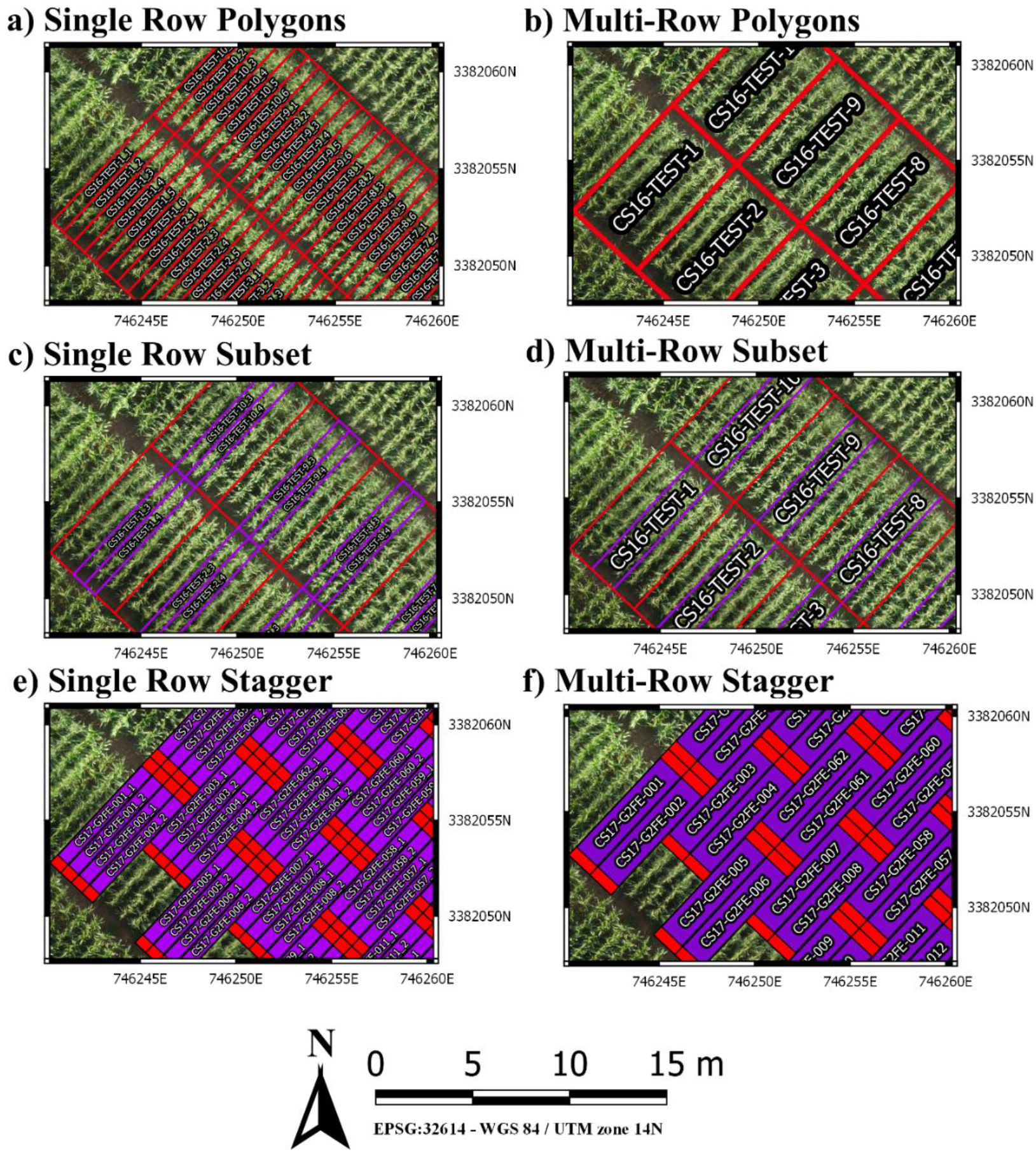
**[a]** Polygons created for each individual row of a six row plot. **[b]** Single polygons created for each plot merging the adjacent rows of each plot. **[c]** Sub-setting out the middle two rows (purple) of a six row plot (red). **[d]** Sub-setting out the middle two rows and merging them to a single polygon (purple) of a six row plot (red). **[e]** Staggering individual row plot polygons to adjust for staggered planting. **[f]** Staggering merged two row plot polygons to adjust for staggered planting.

It is common in advance trials to collect data from interior rows of a multi-row plot to factor out neighbor plot competition. Plotshpcreate has a built in sub setting functionally (plotsubset=”n”) to create polygons of those specific AOIs. The plotsubset argument works by removing “n” rows from either side of the multi-row plot and returns the remaining inter rows and can be used in combination with “multirowind” and “nrowplot” arguments (Figure 2c and d). For example, with a 6-row plot set “plotsubset=2”, plotshpcreate will return the inner 2 rows of the plot removing 2 rows from adjacent sides. Alternatively, all 6 individual plots could be extracted and the outer 4 discarded, however this would result in a threefold larger file taking additional time to extract and analyze, and the two inner rows would still need to be averaged in some appropriate way.

Furthermore, plotshpcreate can adjust polygon geolocation based on a consistent staggering of plot plantings caused by GIS or tripping issues. Plotshpcreate can create staggered plots grids with an impute vector describing the row where staggers start, how many rows the planter sows in a pass, and the stagger offset. For example, if we set “stagger=c(5,4,3.8)”, plotshpcreate will create a four row stagger, 3.8 m towards the back of the field based on the “A” point, beginning at fifth row of experimental design (Figure 2e and f). The functionality of the row which the stagger begins is based on the possibility of border rows, if you have 3 rows of border and a 4 row planter, the stagger would begin on the second row of the trial (e.g. stagger=c(2,4,3.8), if border is not included within your input file.

## 3. Discussion

Implementation of high throughput phenotyping platforms such as UAS or ground vehicles can provide a vast amount of data rapidly. In contrast, the development of tools to process sensor datasets is in its infancy, or non-existent, and continued development of data analytic tools is critical to aid rapid data analysis for actionable information extraction (Shakoor et al., 2019). As a result, manual data wrangling remains a laborious time sink in processing sensor datasets. Plotshpcreate was developed to overcome a critical time sink within the data processing pipeline, creating AOIs for research plots at scale. Plotshpcreate provides a tool to rapidly create gridded AOI polygons with attached unique IDs for extraction of sensor data on an agriculture research plot scale within seconds, compared to the hours it would require to manually draw polygons and define unique IDs of thousands of plots within a GIS software. Foundational tools, like plotshpcreate, set the basis for developing more advanced point and click graphical user interface tools, such as shiny (Chang et al., 2019). Additionally, incorporating algorithms that utilize the imagery to auto correct for minor changes in plot orientation (Ribera et al., 2017) would be a useful, although it would likely increase computation time and memory with the inclusion of imagery data analysis. Plotshpcreate has room for improvement through increased functionality and the developers encourage the community to aid in adding new tools and they feel necessary.

## Supplementary Materials

**File S1**: R/UAStools v0.2.0 R package.

## Author Contribution

Conceptualization, S.A. and S.M.; Methodology, S.A. and S.M..; Software, S.A. and S.M.; Validation, S.A.; Resources, X.X.; Data Curation, S.A.; Writing – Original Draft Preparation, S.A.; Writing – Review & Editing, S.A. and S.M..; Visualization, S.A.; Supervision, S.M.; Project Administration, S.M.; Funding Acquisition, S.M.

## References

Araus, J.L., Kefauver, S.C., Zaman-Allah, M., Olsen, M.S., and Cairns, J.E. (2018). Translating high-throughput phenotyping into genetic gain. Trends in Plant Science 23(5), 451–466. doi:10.1016/j.tplants.2018.02.001.

Bivand, R., Keitt, T., and Rowlingson, B. (2019). rgdal: Bindings for the ’Geospatial’ Data Abstraction Library. R package version 1.4-4. https://CRAN.R-project.org/package=rgdal.

Bivand, R.S., Pebesma, E., and Gomez-Rubio, V. (2013). Applied spatial data analysis with {R}, Second edition. Springer, NY.

Chang, W., Cheng, J., Allaire, J., Xie, Y., and McPherson, J. (2019). shiny: Web Application Framework for R. R package version 1.3.2. https://CRAN.R-project.org/package=shiny.

ESRI Development Team (2019). ArcGIS Pro. Environmental Systems Research Institute (ESRI), http://resources.arcgis.com/en/help/main/10.2/index.html.

Pebesma, E.J., and Bivand, R.S. (2005). Classes and methods for spatial data in R. R News 5(2), 9–13.

Progeny Development Team (2019). Progeny. Progeny Drone Inc. https://www.progenydrone.com/.

QGIS Development Team (2019). QGIS Geographic Information System. Open Source Geospatial Foundation Project http://qgis.osgeo.org.

Ribera, J., Chen, Y., Boomsma, C., and Delp, E.J. (2017). Counting plants using deep learning. 2017 IEEE global conference on signal and information processing (GlobalSIP), 1344–1348. doi:10.1109/GlobalSIP.2017.8309180.

Shakoor, N., Lee, S., and Mockler, T.C. (2017). High throughput phenotyping to accelerate crop breeding and monitoring of diseases in the field. Current opinion in plant biology 38, 184–192. doi:10.1016/j.pbi.2017.05.006.

Shakoor, N., Northrup, D., Murray, S., and Mockler, T.C. (2019). Big Data Driven Agriculture: Big Data Analytics in Plant Breeding, Genomics, and the Use of Remote Sensing Technologies to Advance Crop Productivity. The Plant Phenome Journal 2(1). doi:10.2135/tppj2018.12.0009.

