## Supplemental File 1 for "R/UAStools::plotshpcreate: Create Multi-Polygon Shapefiles for Extraction of Research Plot Scale Agriculture Remote Sensing Data": DESCRIPTION

Package: UAStools
Type: Package
Title: Tools for Field Based Remote Sensing Applications within Plot Based Agriculture 
Version: 0.3.0
Date: 12/09/2019
Author: Steven L. Anderson II,
        Seth C. Murray
Maintainer: Steven L. Anderson II <>
Description: This package provide a function to automate field plot shapefile (.shp) polygons for use in data extraction of remote sensing data sets.
License: GPL-2
Encoding: UTF-8
LazyData: true
Imports:rgdal (>= 1.2-18), sp (>= 1.2-7)
RoxygenNote: 7.0.2
