## Supplemental File 1 for "R/UAStools::plotshpcreate: Create Multi-Polygon Shapefiles for Extraction of Research Plot Scale Agriculture Remote Sensing Data": plotshpcreate.R

```
#' @title Plot Shapefile Create
#'
#' @description Methods for constructing multipolygon ESRI shapefiles (.shp)
#' with individual polygons containing agricultural plot boundaries. Utilizes
#' AB line to rotate polygons to the appropriate geospatial direction of research
#' plots, allowing for simple overlay upon Unmanned Aerial System (UAS) or other
#' geospatial datasets (e.g. orthomosaics, point clouds, etc.) for feature extraction.
#' 
#' It is recommended to define working directory for output files prior to executing plotshpcreate.
#'
#' @param A Vector of UTM coordinates (Easting,Northing) of "A" point, which corresponds
#' to the bottom left corner of the first field plot.
#' Note: This point is where the function begins building polygons. This point is intended
#' to be the midpoint of alley at the front of the first plot and the midpoint of the
#' @param B Vector of UTM coordinates (Easting,Northing) of "B" point, which corresponds
#' to the top left corner of the field trial within the same row as the "A" point
#' @param UTMzone Character parameter defining UTM zone number. Default is NULL and will result in an coordinate reference system of "NA".
#' @param Hemisphere Character parameter that designates the Northern "N" or Southern "S" Hemisphere. Default is "N".
#' @param infile Data frame containing seed preperation file and experimental design
#' (i.e. coordinates of plots within the design grid)
#' @param outfile Character assignment to define outputfile names. If left null, default
#' file names will be produced.
#' @param nrowplot Number of adjacent rows that constitute a plot.Setting this parameter
#' to greater than one allows for different polygon options. Default set to 1.
#' @param multirowind Logic parameter that indicates if adjacent plot rows should be
#' combined and treated as a single plot shapefile and unique idenfifier. Setting to false
#' will combine adjacent plots and construct a single polygon that encompasses
#' all adjacent rows within a plot. Default set to true.
#' @param rowspc Row (i.e. column) spacing of a single row. Default 2.5 feet in reference to
#' 30 inch row spacing.
#' @param rowbuf Distance removed from both sides of rowspc to create a buffer zone between
#' plots boundaries. Default is 0.1 feet.
#' @param rangespc Range (i.e. row) spacing of a single row. This dimesnsion referes to the total
#' plot length including half alley distance on either side of the plot. Default 25 feet.
#' @param rangebuf Distance removed from both sides of rangespc to create a buffer zone between
#' plots boundaries. Default is 2 feet in reference to four foot allies. Note: If alleys are
#' 4 feet rangebuf should be set to 2 feet to remove from both ends of the polygon.
#' @param stagger Numeric vector of length three defining [1] row where staggers starts, 
#' [2] rows sowed by planted in a single pass, and [3] stagger offset distance from A point.
#' @param plotsubset Defines how many adjacent rows should be excluded from the shapefile.
#' This parameter is useful for multirow plots where the central plots are of interest. If plotsubset=1,
#' the outer row will be removed from both sides of the plot.
#' @param field Character vector to indicate the trial the shapefile is being developed for.
#' Recommended format example: CS17-G2FE, which inicated location (CS), year (17), and trial (G2FE).
#' @param unit Character vector that the unit of measure for the polygon dimesnions. Default is "feet".
#' Units can also be input as meters using unit="meter".
#' @param SquarePlot Logic parameter to indicated if PDF file is desired for visualization of none rotated polygons.
#' @param RotatePlot Logic parameter to indicated if PDF file is desired for visualization of rotated polygons.
#' @import rgdal sp``
#' @export
#' @return NULL 
#' @note it is recommendeed to repeat unique Barcodes and Plot numbers if there are multirow plots (mrowplot>1) as
#' the plotshpcreatre function accounts for this redundance within the function.
#'
#' @examples
#'
#'
#' ### Creates shape file for each individual with the inclusion of multirow plot design
#' ### with nrowplot>1 and multirowind=T. Unique identifiers are subset by the location
#' ### of the adjacent plots reading from left to right in geographical space.
#' 
#' ### Set working directory to where you want the files to be saved
#' setwd("C://Temp")
#'
#' plotshpcreate(A=c(746239.817,3382052.264), #Point A c(Easting_0.0,Northing_0.0)
#'               B=c(746334.224,3382152.870), #Point B c(Easting_1.0,Northing_1.0)
#'               UTMzone="14",
#'               Hemisphere="N",
#'               infile=SampleInfile,
#'               outfile="Ind_Multirow_plots",
#'               nrowplot=2,
#'               multirowind=TRUE,
#'               rowspc=2.5,
#'               rowbuf=0.1, ### Will take off both sides of the plot
#'               rangespc=25,
#'               rangebuf=2, ### Half the alley width
#'               plotsubset=NULL,
#'               field="CS17-G2FE",
#'               unit="feet",
#'               SquarePlot=TRUE,
#'               RotatePlot=TRUE)
#'
#' # Creates shape file by combining adacent row of unique plots of multirow plot design
#' # within a single polygone with nrowplot>1 and multirowind=T. If a plot is two rows wide
#' # the shape file will encompass the plot as a whole.
#' 
#' ### Set working directory to where you want the files to be saved
#' setwd("C://Temp")
#' plotshpcreate(A=c(746239.817,3382052.264), #Point A c(Easting_0.0,Northing_0.0)
#'               B=c(746334.224,3382152.870), #Point B c(Easting_1.0,Northing_1.0)
#'               UTMzone="14",
#'               Hemisphere="N"
#'               infile=SampleInfile,
#'               outfile="Multirowplotscombined",
#'               nrowplot=2,
#'               multirowind=FALSE,
#'               rowspc=2.5,
#'               rowbuf=0.1, ### Will take off both sides of the plot
#'               rangespc=25,
#'               rangebuf=2, ### Half the alley width
#'               plotsubset=NULL,
#'               field="CS17-G2FE",
#'               unit="feet",
#'               SquarePlot=TRUE,
#'               RotatePlot=TRUE)
#'
#' # If the experiment is a single row plot design utilize nrowplot=1.


#### x<-read.csv("C://Temp//plotshpcreate_6plottest.csv",header=T)

plotshpcreate<-function(A=NULL, #Point A c(Easting_0.0,Northing_0.0)
                        B=NULL, #Point B c(Easting_1.0,Northing_1.0)
                        UTMzone=NULL,
                        Hemisphere="N",
                        infile=NULL,
                        outfile=NULL,
                        nrowplot=1,
                        multirowind=FALSE,
                        rowspc=2.5,
                        rowbuf=0.1, ### Will take off both sides of the plot
                        rangespc=25,
                        rangebuf=2, ### Half the alley width
                        stagger=NULL, ### c(row where staggers starts, planter rows, stagger distance)
                        plotsubset=0,
                        field=NULL,
                        unit="feet",
                        SquarePlot=TRUE,
                        RotatePlot=TRUE){
# infile<-x
  if (!requireNamespace("rgdal", quietly = TRUE)) {
    stop("Package \"rgdal\" needed for this function to work. Please install it.",
         call. = FALSE)
  }
  
  if (!requireNamespace("sp", quietly = TRUE)) {
    stop("Package \"sp\" needed for this function to work. Please install it.",
         call. = FALSE)
  }
  
  if (!is.null(stagger)){
    if(stagger[[1]]==1){
      stop("Stagger must be in reference to plots beyond first plot polygon, i.e. stagger[[1]]!=1",
           call. = FALSE)
    }
    if(stagger[[1]]>(stagger[[2]]+1)){
      stop("Stagger is based on planter dimesions ans was built for consistent stagger throughout the trial, 
           i.e. stagger[[1]] !> (stagger[[2]]+1)",
           call. = FALSE)
    }
    if(nrowplot>1 & multirowind==F & (nrowplot>(stagger[[2]]/2))){
    stop("Combined plots will not be correctly adjusted by stagger due to:

                   nrowplot>(stagger[[2]]/2)

         ~~~Recommend setting multirowind=T~~~",
         call. = FALSE)
    }
    if(nrowplot>1 & multirowind==F & ((stagger[[2]]%%nrowplot)!=0)){
      stop("Combined plots will not be correctly adjusted by stagger due to:

                          stagger[[2]]%%nrowplot)!=0

            ~~~Recommend setting multirowind=T~~~~",
           call. = FALSE)
    }
    if(nrowplot>1 & multirowind==F & 
       (((stagger[[2]]-(stagger[[2]]-stagger[[1]]-1))%%nrowplot)!=0)){
      stop("Combined plots will not be correctly adjusted by stagger due to:

              (((stagger[[2]]-(stagger[[2]]-stagger[[1]]-1))%%nrowplot)!=0)

            ~~~Recommend setting multirowind=T~~~~",
           call. = FALSE)
    }
  }


  options(digits=12)

  ### Order infile by plot numbers numerically, then by row number 
  infile<-infile[order(as.numeric(infile$Plot),as.numeric(infile$Row)),]

  nRange<- length(unique(infile$Range))
  nRow<- length(unique(infile$Row))

  nPlot<- nrow(infile) 
  
  if (nPlot != (nRange*nRow))
  {warning("Length of file is not equal to number of plots: may cause errors, please remove blank rows at end of file and check row/range numbering")}

  # rm(FirstPlot,LastPlot)
  for (i in 1:nPlot){
    ### Find bottom left plot
    if (infile$Range[i]==min(infile$Range)&& infile$Row[i]==min(infile$Row)) {BLCornerPlot<-i}
    ### Find bottom right plot
    if (infile$Range[i]==min(infile$Range)&& infile$Row[i]==max(infile$Row)) {BRCornerPlot<-i}
    ### Find top right plot
    if (infile$Range[i]==max(infile$Range)&& infile$Row[i]==max(infile$Row)) {TRCornerPlot<-i}
    ### Find back left plot
    if (infile$Range[i]==max(infile$Range)&& infile$Row[i]==min(infile$Row)) {TLCornerPlot<-i}
    }

  ### Takes AB line diffences for delta calculation
  DeltaEasting<- B[[1]]-A[[1]]
  DeltaNorthing<- B[[2]]-A[[2]]

  # DeltaEasting<- Easting_1.0-Easting_0.0
  # DeltaNorthing<- Northing_1.0-Northing_0.0

  #Calculates theta in Radians
  DirectionTheta<- atan(abs(DeltaNorthing)/abs(DeltaEasting)) #Radians

### R circles rotate counter clockwise starting at [0,1] i.e right 90 deg of circle 
### Theta is for plot orientation in radians
### srt_theta is for text rotation in plots in degress
  ## QUADRANT I ##
  if(DeltaNorthing>0 && DeltaEasting>0){
    Theta<-(((3*pi)/2)+DirectionTheta)
    srt_theta<-90-DirectionTheta*(180/pi)
  }
  ## QUADRANT II ##
  if(DeltaNorthing>0 && DeltaEasting<0){
    Theta<-(((pi)/2)-DirectionTheta)
    srt_theta<-270+DirectionTheta*(180/pi)
  }
  ## QUADRANT III ##
  if(DeltaNorthing<0 && DeltaEasting<0){
    Theta<-(((pi)/2)+DirectionTheta)
    srt_theta<-DirectionTheta*(180/pi)
  }
  ## QUADRANT IV ##
  if(DeltaNorthing<0 && DeltaEasting>0){
    Theta<-(((3*pi)/2)-DirectionTheta)
    srt_theta<-270+DirectionTheta*(180/pi)
  }

  if(unit=="feet"){
    RangeSpacingM<-rangespc/3.281 #meter
    RowSpacingM<- rowspc/3.281 #meter
    RangeBufferM<-rangebuf/3.281 #meter
    RowBufferM<- rowbuf/3.281 #meter
    staggerM<-stagger[[3]]/3.281
    }

  if(unit=="meter"){
    RangeSpacingM<-rangespc #meter
    RowSpacingM<- rowspc #meter
    RangeBufferM<-rangebuf #meter
    RowBufferM<- rowbuf #meter
    staggerM<-stagger[[3]]
    }


  ZeroZero<-matrix(0,1,2)
  ZeroZero[1,1] <- 0 #Easting_0.0
  ZeroZero[1,2] <- 0 #Northing_0.0


  ### When plots are perfectly perpendicular as if planted perfectly north to south
  ### column 12 added to retain orginial row numbering for stagger
  PlotsSquareM<-matrix(NA,nPlot,12)
  PlotsSquareM[,1]<- infile$Plot # Plot number
  PlotsSquareM[,2]<- infile$Range-min(infile$Range)+1 #RANGE = Y
  PlotsSquareM[,3]<-PlotsSquareM[,12]<- infile$Row-min(infile$Row)+1 #ROW = X
  rownames(PlotsSquareM)<-rep(NA,nPlot)
  
  if(nrowplot==1){
    rownames(PlotsSquareM)<-infile$Barcode
  }
  # PlotsSquareM[,12]<- as.character(infile$Barcode) # Plot number

  ####################################################################
  ####################################################################
  ##### Combining Multi-row adjacent plots to a single plot ##########
  ####################################################################
  ####################################################################
  if(nrowplot>1 & multirowind==F & plotsubset==0){

    if(unit=="feet"){
      RangeSpacingM<-rangespc/3.281 #meter
      RowSpacingM<- (nrowplot*rowspc)/3.281 #meter
      RangeBufferM<-rangebuf/3.281 #meter
      RowBufferM<- rowbuf/3.281 #meter
      staggerM<-stagger[[3]]/3.281
    }

    if(unit=="meter"){
      RangeSpacingM<-rangespc #meter
      RowSpacingM<- (nrowplot*rowspc) #meter
      RangeBufferM<-rangebuf #meter
      RowBufferM<- rowbuf #meter
      staggerM<-stagger[[3]]
    }

    # rowspc<-rowspc*nrowplot

    ID<-unique(as.character(infile$Barcode))

    for(i in 1:length(ID)){
      
      #### Finds rows in PlotsSquareM 
      rownames(PlotsSquareM)[which(PlotsSquareM[,1]==infile$Plot[infile$Barcode==ID[[i]]])]<-ID[i]
    }

    PlotsSquareM<-PlotsSquareM[PlotsSquareM[,3]==(1+plotsubset) | PlotsSquareM[,3] %in% seq(1+nrowplot+plotsubset,max(PlotsSquareM[,3]),nrowplot),]

    # nPlot<-nrow(PlotsSquareM)

    PlotsSquareM[,12]<-PlotsSquareM[,3]
    for (i in 1:length(PlotsSquareM[,3])){
      if(PlotsSquareM[i,3]==1){next}
      else(PlotsSquareM[i,3]<-ceiling(PlotsSquareM[i,3]/nrowplot))
    }


    ########### Subseting out region of larger plot ################
    if(plotsubset!=0){
      if(nrowplot==1){
        stop("nrowplot == 1 : Cannot subset singular plot.",
                       call. = FALSE)
        }
      if(nrowplot<3){
        stop("nrowplot < 3 : Cannot subset central plot. \n Recommend nrowplot=2, multirowind==T,plotsubset=NULL. ",
             call. = FALSE)
      }
      if(nrowplot==2*plotsubset){
        stop("nrowplot==2*plotsubset : no polygons will be created ",
             call. = FALSE)
      }

    }

      rowbuff<-rowbuf+(rowspc*plotsubset)
}

  ####################################################################
  ####################################################################
  ##### Formatting Adjacent Plot to Unique IDs w/o combining #########
  ####################################################################
  ####################################################################
  if((nrowplot>1 & multirowind==T)|(plotsubset!=0)){

    IDs<-unique(as.character(infile$Barcode))

  PolygonsToMake <- vector('list', length(IDs))

  # i=1
  # k=1
  for(i in 1:length(IDs)){
    # for (j in 1:nRow){
    ### Subsets out all rows of  PlotsSquareM whwere the row numbers are 
    ### equal to the unique barcode for each iteration if "i"
    
      subset_PSM<-PlotsSquareM[which(PlotsSquareM[,1]==infile$Plot[infile$Barcode==IDs[[i]]]),]

      # subset_PSM<-rbind(subset_PSM,subset_PSM)
      # subset_PSM[2,3]<-50
      nrow_subset<-nrow(subset_PSM)
      for (k in 1:nrow_subset){
        
      ### Looking for the min row number of plots with the same plot number
      ### iteritively creating unquie barcodes with left to right numbering of rows 1 to nrowplot
        if(k==nrow_subset){

          rownames(PlotsSquareM)[which(PlotsSquareM[,1]==subset_PSM[1] &
                                         PlotsSquareM[,3]==subset_PSM[3])]<-paste(IDs[[i]],k,sep="_")

        }

        if(k<nrow_subset){
        rownames(PlotsSquareM)[which(PlotsSquareM[,1]==subset_PSM[subset_PSM[,3]==min(subset_PSM[,3]),1] &
                                       PlotsSquareM[,3]==subset_PSM[subset_PSM[,3]==min(subset_PSM[,3]),3])]<-paste(IDs[i],k,sep="_")

        subset_PSM<-subset_PSM[-which(subset_PSM[,3]==min(subset_PSM[,3])),]
        }

      }
  }
  #### print(PlotsSquareM)

  if(plotsubset!=0){
    if(nrowplot==1){
      stop("nrowplot == 1 : Cannot subset singular plot.",
           call. = FALSE)
    }
    if(nrowplot<3){
      stop("nrowplot < 3 : Cannot subset central plot. \n Recommend nrowplot=2, multirowind==T,plotsubset=0. ",
           call. = FALSE)
    }
    if(nrowplot==2*plotsubset){
      stop("nrowplot==2*plotsubset : no polygons will be created ",
           call. = FALSE)
    }
    
  ### PlotsSquareM[,13] is a logical column to identify the rows to include if plotsubset is desired
  PlotsSquareM<-cbind(PlotsSquareM,rep(F,nrow(PlotsSquareM)))

  ### This loop identifies the interior plots of the multirowplot to retain after subset  
for (i in 1:nrow(PlotsSquareM)){
      PlotsSquareM[i,13]<-PlotsSquareM[i,3] %in% 
        seq(
          min(PlotsSquareM[PlotsSquareM[,1]==PlotsSquareM[i,1],3])+plotsubset,
          max(PlotsSquareM[PlotsSquareM[,1]==PlotsSquareM[i,1],3])-plotsubset,
          1)
}
  # PlotsSquareM<-PlotsSquareM[PlotsSquareM[,13]==1,]
  }
  }


  ####################################################################
  ####################################################################
  ################ Serpentine ordering of plots ######################
  ####################################################################
  ####################################################################
  for (i in 1:nRange){

    subset_PSM<-PlotsSquareM[PlotsSquareM[,2]==i,]

    if (i%%2==0){
      subset_PSM<-subset_PSM[order(subset_PSM[,3],decreasing=T),]
    }

    if (i%%2!=0){
      subset_PSM<-subset_PSM[order(subset_PSM[,3],decreasing=F),]
    }

    if(i==1){
      new_PSM<-subset_PSM
    }

    if(i!=1){
      new_PSM<-rbind(new_PSM,subset_PSM)
    }
  }

  PlotsSquareM<-PlotSquare4plot<-new_PSM
  
  ### Subsets out the interior plots when multirowind==T
  if(nrowplot>1 & multirowind==T & plotsubset!=0){
    PlotsSquareM<-PlotSquare4plot<-new_PSM[new_PSM[,13]==1,]
  }

  ####################################################################
  ####################################################################
  ####################################################################
  ####################################################################
  ### remember the point is in the middle of the plot not the furrow
  #### ^^^^^ I Think this is wrong -Steve 1/22/2019

  # i<-2
  
  # PlotsSquareM<-PlotsSquareM[PlotsSquareM[,13]==1,]
  
  for (i in 1:nrow(PlotsSquareM)) {
   
      
      PlotsSquareM[i,4]<- ZeroZero[1,1]+((PlotsSquareM[i,2]-1)*RangeSpacingM)  # Y bottom left Range
      PlotsSquareM[i,5]<- ZeroZero[1,2]+((PlotsSquareM[i,3]-1)* RowSpacingM)   # X bottom left Row
      
      PlotsSquareM[i,6]<- ZeroZero[1,1]+((PlotsSquareM[i,2]-1)*RangeSpacingM )   # Y bottom right Range
      PlotsSquareM[i,7]<- ZeroZero[1,2]+((PlotsSquareM[i,3])* RowSpacingM)   # X bottom right Row
      
      PlotsSquareM[i,8]<- ZeroZero[1,1]+((PlotsSquareM[i,2])*RangeSpacingM )  # Y top right Range
      PlotsSquareM[i,9]<- ZeroZero[1,2]+((PlotsSquareM[i,3])* RowSpacingM)   # X top right Row
      
      PlotsSquareM[i,10]<- ZeroZero[1,1]+((PlotsSquareM[i,2])*RangeSpacingM )  # Y top left Range
      PlotsSquareM[i,11]<- ZeroZero[1,2]+((PlotsSquareM[i,3]-1)* RowSpacingM)  # X top left Row
    
     
    
    
    if(!is.null(stagger)){
      # 
      # 
      # if (!is.null(stagger)){
      #   stag_vector<-c()
      #   for (i in 1:nrow(PlotsSquareM)){
      #     if(PlotsSquareM[i,3]<stagger[[1]]){
      #       stag_vector[i]<-0
      #       next
      #     }
      #     if(PlotsSquareM[i,3]==stagger[[1]]){
      #         stag_vector[i]<-1
      #         next}
      #     if(PlotsSquareM[i,3]>=stagger[[1]]){
      #       a<-floor((PlotsSquareM[i,3]-stagger[[1]])+1)
      #       b<-ceiling((a+stagger[[2]])/stagger[[2]])
      #       if(b%%2==0
      #     }
      # 
      #       }
      #     }
      #   }
      #   stag_vector<-rep(c(rep(1,stagger[[2]]),rep(0,stagger[[2]])),(nRow-stagger[[1]])/(stagger[[2]]*2))

      
      
      if((ceiling((floor((PlotsSquareM[i,12]-stagger[[1]])+1)+stagger[[2]])/stagger[[2]])%%2)==0){
    
    PlotSquare4plot[i,4]<- ZeroZero[1,1]+((PlotsSquareM[i,2]-1)*RangeSpacingM)+staggerM  # Y bottom left Range
    PlotSquare4plot[i,5]<- ZeroZero[1,2]+((PlotsSquareM[i,3]-1)* RowSpacingM)   # X bottom left Row

    PlotSquare4plot[i,6]<- ZeroZero[1,1]+((PlotsSquareM[i,2]-1)*RangeSpacingM )+staggerM   # Y bottom right Range
    PlotSquare4plot[i,7]<- ZeroZero[1,2]+((PlotsSquareM[i,3])* RowSpacingM)   # X bottom right Row

    PlotSquare4plot[i,8]<- ZeroZero[1,1]+((PlotsSquareM[i,2])*RangeSpacingM )+staggerM   # Y top right Range
    PlotSquare4plot[i,9]<- ZeroZero[1,2]+((PlotsSquareM[i,3])* RowSpacingM)   # X top right Row

    PlotSquare4plot[i,10]<- ZeroZero[1,1]+((PlotsSquareM[i,2])*RangeSpacingM )+staggerM   # Y top left Range
    PlotSquare4plot[i,11]<- ZeroZero[1,2]+((PlotsSquareM[i,3]-1)* RowSpacingM)  # X top left Row
  }
      
      if((ceiling((floor((PlotsSquareM[i,12]-stagger[[1]])+1)+stagger[[2]])/stagger[[2]])%%2)!=0){
        
        PlotSquare4plot[i,4]<- ZeroZero[1,1]+((PlotsSquareM[i,2]-1)*RangeSpacingM)  # Y bottom left Range
        PlotSquare4plot[i,5]<- ZeroZero[1,2]+((PlotsSquareM[i,3]-1)* RowSpacingM)   # X bottom left Row
        
        PlotSquare4plot[i,6]<- ZeroZero[1,1]+((PlotsSquareM[i,2]-1)*RangeSpacingM )   # Y bottom right Range
        PlotSquare4plot[i,7]<- ZeroZero[1,2]+((PlotsSquareM[i,3])* RowSpacingM)   # X bottom right Row
        
        PlotSquare4plot[i,8]<- ZeroZero[1,1]+((PlotsSquareM[i,2])*RangeSpacingM )   # Y top right Range
        PlotSquare4plot[i,9]<- ZeroZero[1,2]+((PlotsSquareM[i,3])* RowSpacingM)   # X top right Row
        
        PlotSquare4plot[i,10]<- ZeroZero[1,1]+((PlotsSquareM[i,2])*RangeSpacingM )   # Y top left Range
        PlotSquare4plot[i,11]<- ZeroZero[1,2]+((PlotsSquareM[i,3]-1)* RowSpacingM)  # X top left Row
      }
    }
    }
    

  #PlotsSquareM[c(BLCornerPlot, BRCornerPlot, TRCornerPlot, TLCornerPlot, FirstPlot, LastPlot),]

  ### THis is the red boxes
  ## has the data we care about
  
  
  PlotsSquareMBuf<-PlotsSquareBuf4plot<-PlotsSquareM
  
  for (i in 1:nrow(PlotsSquareMBuf)) {
    
    
      
      PlotsSquareMBuf[i,4]<- ZeroZero[1,1]+((PlotsSquareMBuf[i,2]-1)*RangeSpacingM ) + RangeBufferM  # Y bottom left
      PlotsSquareMBuf[i,5]<- ZeroZero[1,2]+((PlotsSquareMBuf[i,3]-1)* RowSpacingM) + RowBufferM # X bottom left
      
      PlotsSquareMBuf[i,6]<- ZeroZero[1,1]+((PlotsSquareMBuf[i,2]-1)*RangeSpacingM ) +RangeBufferM # Y bottom right
      PlotsSquareMBuf[i,7]<- ZeroZero[1,2]+((PlotsSquareMBuf[i,3])* RowSpacingM) - RowBufferM  # X bottom right
      
      PlotsSquareMBuf[i,8]<- ZeroZero[1,1]+((PlotsSquareMBuf[i,2])*RangeSpacingM ) -RangeBufferM # Y top right
      PlotsSquareMBuf[i,9]<- ZeroZero[1,2]+((PlotsSquareMBuf[i,3])* RowSpacingM) - RowBufferM # X top right
      
      PlotsSquareMBuf[i,10]<- ZeroZero[1,1]+((PlotsSquareMBuf[i,2])*RangeSpacingM ) - RangeBufferM  # Y top left
      PlotsSquareMBuf[i,11]<- ZeroZero[1,2]+((PlotsSquareMBuf[i,3]-1)* RowSpacingM) + RowBufferM # X top left
  
    
        
    if(!is.null(stagger)){
      if((ceiling((floor((PlotsSquareM[i,12]-stagger[[1]])+1)+stagger[[2]])/stagger[[2]])%%2)==0){
        
        PlotsSquareBuf4plot[i,4]<- ZeroZero[1,1]+((PlotsSquareMBuf[i,2]-1)*RangeSpacingM ) + RangeBufferM +staggerM   # Y bottom left
        PlotsSquareBuf4plot[i,5]<- ZeroZero[1,2]+((PlotsSquareMBuf[i,3]-1)* RowSpacingM) + RowBufferM # X bottom left

        PlotsSquareBuf4plot[i,6]<- ZeroZero[1,1]+((PlotsSquareMBuf[i,2]-1)*RangeSpacingM ) +RangeBufferM +staggerM  # Y bottom right
        PlotsSquareBuf4plot[i,7]<- ZeroZero[1,2]+((PlotsSquareMBuf[i,3])* RowSpacingM) - RowBufferM  # X bottom right

        PlotsSquareBuf4plot[i,8]<- ZeroZero[1,1]+((PlotsSquareMBuf[i,2])*RangeSpacingM ) -RangeBufferM +staggerM  # Y top right
        PlotsSquareBuf4plot[i,9]<- ZeroZero[1,2]+((PlotsSquareMBuf[i,3])* RowSpacingM) - RowBufferM # X top right

        PlotsSquareBuf4plot[i,10]<- ZeroZero[1,1]+((PlotsSquareMBuf[i,2])*RangeSpacingM ) - RangeBufferM + staggerM  # Y top left
        PlotsSquareBuf4plot[i,11]<- ZeroZero[1,2]+((PlotsSquareMBuf[i,3]-1)* RowSpacingM) + RowBufferM # X top left
      }
      
    if((ceiling((floor((PlotsSquareM[i,12]-stagger[[1]])+1)+stagger[[2]])/stagger[[2]])%%2)!=0){
      PlotsSquareBuf4plot[i,4]<- ZeroZero[1,1]+((PlotsSquareMBuf[i,2]-1)*RangeSpacingM ) + RangeBufferM  # Y bottom left
      PlotsSquareBuf4plot[i,5]<- ZeroZero[1,2]+((PlotsSquareMBuf[i,3]-1)* RowSpacingM) + RowBufferM # X bottom left
      
      PlotsSquareBuf4plot[i,6]<- ZeroZero[1,1]+((PlotsSquareMBuf[i,2]-1)*RangeSpacingM ) +RangeBufferM # Y bottom right
      PlotsSquareBuf4plot[i,7]<- ZeroZero[1,2]+((PlotsSquareMBuf[i,3])* RowSpacingM) - RowBufferM  # X bottom right
      
      PlotsSquareBuf4plot[i,8]<- ZeroZero[1,1]+((PlotsSquareMBuf[i,2])*RangeSpacingM ) -RangeBufferM # Y top right
      PlotsSquareBuf4plot[i,9]<- ZeroZero[1,2]+((PlotsSquareMBuf[i,3])* RowSpacingM) - RowBufferM # X top right
      
      PlotsSquareBuf4plot[i,10]<- ZeroZero[1,1]+((PlotsSquareMBuf[i,2])*RangeSpacingM ) - RangeBufferM  # Y top left
      PlotsSquareBuf4plot[i,11]<- ZeroZero[1,2]+((PlotsSquareMBuf[i,3]-1)* RowSpacingM) + RowBufferM # X top left
    }
    }
}

      
  if(nrowplot>1 & multirowind==F & plotsubset!=0){
    
    ID<-unique(as.character(infile$Barcode))
    plots<-unique(as.numeric(PlotsSquareM[,1]))
    
    newPlotDF<-data.frame()
    newPlotBufDF<-data.frame()
    for (i in 1:length(plots)){
      pulls<-PlotsSquareM[PlotsSquareM[,1]==plots[i],]
      pulls<-as.data.frame(pulls[pulls[,13]==1,])
      
      pulls1<-pulls[pulls[,12]==min(pulls[,12]),]
      pulls1[,4]<-min(pulls[,4]) # Y bottom left Range
      pulls1[,5]<-min(pulls[,5]) # X bottom left Row
      pulls1[,6]<-min(pulls[,6]) # Y bottom right Range
      pulls1[,7]<-max(pulls[,7]) # X bottom right Row
      pulls1[,8]<-max(pulls[,8]) # Y top right Range
      pulls1[,9]<-max(pulls[,9]) # X top right Row
      pulls1[,10]<-max(pulls[,10]) # Y top left Range
      pulls1[,11]<-min(pulls[,11]) # X top left Row
      rownames(pulls1)<-strsplit(rownames(pulls1),"_")[[1]][1]
      
      newPlotDF<-rbind(newPlotDF,pulls1)
      
      Bpulls<-PlotsSquareMBuf[PlotsSquareMBuf[,1]==plots[i],]
      Bpulls<-as.data.frame(Bpulls[Bpulls[,13]==1,])
      
      Bpulls1<-Bpulls[Bpulls[,12]==min(Bpulls[,12]),]
      Bpulls1[,4]<-min(Bpulls[,4]) # Y bottom left Range
      Bpulls1[,5]<-min(Bpulls[,5]) # X bottom left Row
      Bpulls1[,6]<-min(Bpulls[,6]) # Y bottom right Range
      Bpulls1[,7]<-max(Bpulls[,7]) # X bottom right Row
      Bpulls1[,8]<-max(Bpulls[,8]) # Y top right Range
      Bpulls1[,9]<-max(Bpulls[,9]) # X top right Row
      Bpulls1[,10]<-max(Bpulls[,10]) # Y top left Range
      Bpulls1[,11]<-min(Bpulls[,11]) # X top left Row
      rownames(Bpulls1)<-strsplit(rownames(Bpulls1),"_")[[1]][1]
      
      newPlotBufDF<-rbind(newPlotBufDF,Bpulls1)
    }
    PlotsSquareM<-as.matrix(newPlotDF)
    PlotsSquareMBuf<-as.matrix(newPlotBufDF)
  }
      
  # if(nrowplot>1 & multirowind==F){ 
  #   # for(i in 1:length(ID)){
  #   #   rownames(PlotsSquareM)[which(PlotsSquareM[,1]==infile$Plot[infile$Barcode==ID[[i]]])]<-ID[i]
  #   # }
  #   # 
  #   # PlotsSquareM<-PlotsSquareM[PlotsSquareM[,3]==(1+plotsubset) | PlotsSquareM[,3] %in% seq(1+nrowplot+plotsubset,max(PlotsSquareM[,3]),nrowplot),]
  #   # 
  #   # nPlot<-nrow(PlotsSquareM)
  #   
  #   # PlotsSquareM[,12]<-PlotsSquareM[,3]
  #   for (i in 1:length(PlotsSquareM[,3])){
  #     if(PlotsSquareM[i,3]==1){next}
  #     else(PlotsSquareM[i,3]<-ceiling(PlotsSquareM[i,3]/nrowplot))
  #   }
  #   
  # }
    
  
  ###### PLOTTING SIZE OF PLOT DIMENSIONS ############
  if (SquarePlot==T){
    if(is.null(stagger)){
      PlotSquare4plot<-PlotsSquareM
      PlotsSquareBuf4plot<-PlotsSquareMBuf
    }
    
  PlotY<- cbind(PlotSquare4plot[,4], PlotSquare4plot[,6], PlotSquare4plot[,8], PlotSquare4plot[,10]) #Ranges
  PlotX<- cbind(PlotSquare4plot[,5], PlotSquare4plot[,7], PlotSquare4plot[,9], PlotSquare4plot[,11]) #Rows outline

  PlotYBuf<- cbind(PlotsSquareBuf4plot[,4], PlotsSquareBuf4plot[,6], PlotsSquareBuf4plot[,8], PlotsSquareBuf4plot[,10]) #Ranges buffered
  PlotXBuf<- cbind(PlotsSquareBuf4plot[,5], PlotsSquareBuf4plot[,7], PlotsSquareBuf4plot[,9], PlotsSquareBuf4plot[,11]) #Ranges buffered


  ########## Square Visualization of Trial Plots ##############
  pdf(paste(field,outfile,"Square_plots.pdf",sep="_"),
      width=15,
      height=15)
  
  plot(c(0,nRow*RowSpacingM), c(0,nRange*RangeSpacingM), type="n", xlab = "ROWS (meters)", ylab = "RANGES (meters)") # Plot lines in field
  
  if(nrowplot>1 & multirowind==F & plotsubset==0){
  plot(c(0,max(PlotSquare4plot[,3])*RowSpacingM), c(0,nRange*RangeSpacingM), type="n", xlab = "ROWS (meters)", ylab = "RANGES (meters)") # Plot lines in field
    }
    
  for (i in 1:nrow(PlotSquare4plot)) {
    for (j in 1:4) {
      points(PlotX[i,j], PlotY[i,j])
    }
    polygon(PlotX[i,], PlotY[i,])
    polygon(PlotXBuf[i,], PlotYBuf[i,], col="RED")
    text( ((PlotXBuf[i,1]+PlotXBuf[i,3])/2), ((PlotYBuf[i,1]+PlotYBuf[i,3])/2), labels= PlotsSquareM[i,1], pos = NULL, col = "WHITE", cex=0.7,srt=0 )
  }
  dev.off()
  }


  ########## ADJUST PLOT CORNERS TO NOT CENTER OF PLOT ##############

  ThetaPlotCorner<-Theta
  EastingCorner_0.0<-A[1]
  NorthingCorner_0.0<-A[2]

  if(!is.null(stagger)){
    ## QUADRANT I ##
  if(DeltaNorthing>0 && DeltaEasting>0){
    EastingCorner_0.0_stag<-EastingCorner_0.0-staggerM*sin(ThetaPlotCorner)
    NorthingCorner_0.0_stag<-NorthingCorner_0.0+staggerM*cos(ThetaPlotCorner)
  }
  ## QUADRANT II ##
  if(DeltaNorthing>0 && DeltaEasting<0){
    EastingCorner_0.0_stag<-EastingCorner_0.0-staggerM*cos(ThetaPlotCorner)
    NorthingCorner_0.0_stag<-NorthingCorner_0.0+staggerM*sin(ThetaPlotCorner)
  }
  ## QUADRANT III ##
  if(DeltaNorthing<0 && DeltaEasting<0){
    EastingCorner_0.0_stag<-EastingCorner_0.0-staggerM*cos(ThetaPlotCorner)
    NorthingCorner_0.0_stag<-NorthingCorner_0.0+staggerM*sin(ThetaPlotCorner)
  }
  ## QUADRANT IV ##
  if(DeltaNorthing<0 && DeltaEasting>0){
    EastingCorner_0.0_stag<-EastingCorner_0.0+staggerM*cos(ThetaPlotCorner)
    NorthingCorner_0.0_stag<-NorthingCorner_0.0-staggerM*sin(ThetaPlotCorner)
  }
  }
  
  PlotsAdjustedM<- PlotsSquareM
  PlotsAdjustedMBuf<- PlotsSquareMBuf


  ####################################################################
  ####################################################################
  ######## Rotating Plot Boundaries Based on AB Line #################
  ####################################################################
  ####################################################################
  for (i in 1:nrow(PlotsAdjustedM)) {
    if(is.null(stagger)){
      
      PlotsAdjustedM[i,4]<- (PlotsSquareM[i,4]*cos(Theta))+ (PlotsSquareM[i,5]*sin(Theta)) + NorthingCorner_0.0  # Y bottom left ranges
      PlotsAdjustedM[i,5]<- (PlotsSquareM[i,5]*cos(Theta))- (PlotsSquareM[i,4]*sin(Theta)) + EastingCorner_0.0  # X bottom left rows
      
      #BottomRight
      PlotsAdjustedM[i,6]<- (PlotsSquareM[i,6]*cos(Theta))+ (PlotsSquareM[i,7]*sin(Theta)) + NorthingCorner_0.0   # Y bottom right ranges
      PlotsAdjustedM[i,7]<- (PlotsSquareM[i,7]*cos(Theta))- (PlotsSquareM[i,6]*sin(Theta)) + EastingCorner_0.0  # X bottom right rows
      
      #TopRight
      PlotsAdjustedM[i,8]<- (PlotsSquareM[i,8]*cos(Theta))+ (PlotsSquareM[i,9]*sin(Theta)) + NorthingCorner_0.0   # Y top right ranges
      PlotsAdjustedM[i,9]<- (PlotsSquareM[i,9]*cos(Theta))- (PlotsSquareM[i,8]*sin(Theta)) + EastingCorner_0.0  # X top right rows
      
      #TopLeft
      PlotsAdjustedM[i,10]<- (PlotsSquareM[i,10]*cos(Theta))+ (PlotsSquareM[i,11]*sin(Theta)) + NorthingCorner_0.0   # Y top left ranges
      PlotsAdjustedM[i,11]<- (PlotsSquareM[i,11]*cos(Theta))- (PlotsSquareM[i,10]*sin(Theta)) + EastingCorner_0.0  # X top left rows
      
      ##BottomLeft Adjusted Buffer
      PlotsAdjustedMBuf[i,4]<- (PlotsSquareMBuf[i,4]*cos(Theta))+ (PlotsSquareMBuf[i,5]*sin(Theta)) + NorthingCorner_0.0 # Y bottom left ranges
      PlotsAdjustedMBuf[i,5]<- (PlotsSquareMBuf[i,5]*cos(Theta))- (PlotsSquareMBuf[i,4]*sin(Theta)) + EastingCorner_0.0  # X bottom left rows
      
      #BottomRight Adjusted Buffer
      PlotsAdjustedMBuf[i,6]<- (PlotsSquareMBuf[i,6]*cos(Theta))+ (PlotsSquareMBuf[i,7]*sin(Theta)) + NorthingCorner_0.0# Y bottom right ranges
      PlotsAdjustedMBuf[i,7]<- (PlotsSquareMBuf[i,7]*cos(Theta))- (PlotsSquareMBuf[i,6]*sin(Theta)) + EastingCorner_0.0 # X bottom right rows
      
      #TopRight Adjusted Buffer
      PlotsAdjustedMBuf[i,8]<- (PlotsSquareMBuf[i,8]*cos(Theta))+ (PlotsSquareMBuf[i,9]*sin(Theta)) + NorthingCorner_0.0# Y top right ranges
      PlotsAdjustedMBuf[i,9]<- (PlotsSquareMBuf[i,9]*cos(Theta))- (PlotsSquareMBuf[i,8]*sin(Theta)) + EastingCorner_0.0  # X top right rows
      
      #TopLeft Adjusted Buffer
      PlotsAdjustedMBuf[i,10]<- (PlotsSquareMBuf[i,10]*cos(Theta))+ (PlotsSquareMBuf[i,11]*sin(Theta)) + NorthingCorner_0.0# Y top left ranges
      PlotsAdjustedMBuf[i,11]<- (PlotsSquareMBuf[i,11]*cos(Theta))- (PlotsSquareMBuf[i,10]*sin(Theta)) + EastingCorner_0.0  # X top left rows
    }
    
    
    if(!is.null(stagger)){
      if((ceiling((floor((PlotsSquareM[i,12]-stagger[[1]])+1)+stagger[[2]])/stagger[[2]])%%2)==0){
      
      #BottomLeft
      PlotsAdjustedM[i,4]<- (PlotsSquareM[i,4]*cos(Theta))+ (PlotsSquareM[i,5]*sin(Theta)) + NorthingCorner_0.0_stag  # Y bottom left ranges
      PlotsAdjustedM[i,5]<- (PlotsSquareM[i,5]*cos(Theta))- (PlotsSquareM[i,4]*sin(Theta)) + EastingCorner_0.0_stag  # X bottom left rows
      
      #BottomRight
      PlotsAdjustedM[i,6]<- (PlotsSquareM[i,6]*cos(Theta))+ (PlotsSquareM[i,7]*sin(Theta)) + NorthingCorner_0.0_stag   # Y bottom right ranges
      PlotsAdjustedM[i,7]<- (PlotsSquareM[i,7]*cos(Theta))- (PlotsSquareM[i,6]*sin(Theta)) + EastingCorner_0.0_stag  # X bottom right rows
      
      #TopRight
      PlotsAdjustedM[i,8]<- (PlotsSquareM[i,8]*cos(Theta))+ (PlotsSquareM[i,9]*sin(Theta)) + NorthingCorner_0.0_stag   # Y top right ranges
      PlotsAdjustedM[i,9]<- (PlotsSquareM[i,9]*cos(Theta))- (PlotsSquareM[i,8]*sin(Theta)) + EastingCorner_0.0_stag  # X top right rows
      
      #TopLeft
      PlotsAdjustedM[i,10]<- (PlotsSquareM[i,10]*cos(Theta))+ (PlotsSquareM[i,11]*sin(Theta)) + NorthingCorner_0.0_stag   # Y top left ranges
      PlotsAdjustedM[i,11]<- (PlotsSquareM[i,11]*cos(Theta))- (PlotsSquareM[i,10]*sin(Theta)) + EastingCorner_0.0_stag  # X top left rows
      
      ##BottomLeft Adjusted Buffer
      PlotsAdjustedMBuf[i,4]<- (PlotsSquareMBuf[i,4]*cos(Theta))+ (PlotsSquareMBuf[i,5]*sin(Theta)) + NorthingCorner_0.0_stag # Y bottom left ranges
      PlotsAdjustedMBuf[i,5]<- (PlotsSquareMBuf[i,5]*cos(Theta))- (PlotsSquareMBuf[i,4]*sin(Theta)) + EastingCorner_0.0_stag  # X bottom left rows
      
      #BottomRight Adjusted Buffer
      PlotsAdjustedMBuf[i,6]<- (PlotsSquareMBuf[i,6]*cos(Theta))+ (PlotsSquareMBuf[i,7]*sin(Theta)) + NorthingCorner_0.0_stag# Y bottom right ranges
      PlotsAdjustedMBuf[i,7]<- (PlotsSquareMBuf[i,7]*cos(Theta))- (PlotsSquareMBuf[i,6]*sin(Theta)) + EastingCorner_0.0_stag # X bottom right rows
      
      #TopRight Adjusted Buffer
      PlotsAdjustedMBuf[i,8]<- (PlotsSquareMBuf[i,8]*cos(Theta))+ (PlotsSquareMBuf[i,9]*sin(Theta)) + NorthingCorner_0.0_stag# Y top right ranges
      PlotsAdjustedMBuf[i,9]<- (PlotsSquareMBuf[i,9]*cos(Theta))- (PlotsSquareMBuf[i,8]*sin(Theta)) + EastingCorner_0.0_stag  # X top right rows
      
      #TopLeft Adjusted Buffer
      PlotsAdjustedMBuf[i,10]<- (PlotsSquareMBuf[i,10]*cos(Theta))+ (PlotsSquareMBuf[i,11]*sin(Theta)) + NorthingCorner_0.0_stag# Y top left ranges
      PlotsAdjustedMBuf[i,11]<- (PlotsSquareMBuf[i,11]*cos(Theta))- (PlotsSquareMBuf[i,10]*sin(Theta)) + EastingCorner_0.0_stag  # X top left rows
      
    }
      
      
      if((ceiling((floor((PlotsSquareM[i,12]-stagger[[1]])+1)+stagger[[2]])/stagger[[2]])%%2)!=0){ 
    #BottomLeft
    PlotsAdjustedM[i,4]<- (PlotsSquareM[i,4]*cos(Theta))+ (PlotsSquareM[i,5]*sin(Theta)) + NorthingCorner_0.0  # Y bottom left ranges
    PlotsAdjustedM[i,5]<- (PlotsSquareM[i,5]*cos(Theta))- (PlotsSquareM[i,4]*sin(Theta)) + EastingCorner_0.0  # X bottom left rows

    #BottomRight
    PlotsAdjustedM[i,6]<- (PlotsSquareM[i,6]*cos(Theta))+ (PlotsSquareM[i,7]*sin(Theta)) + NorthingCorner_0.0   # Y bottom right ranges
    PlotsAdjustedM[i,7]<- (PlotsSquareM[i,7]*cos(Theta))- (PlotsSquareM[i,6]*sin(Theta)) + EastingCorner_0.0  # X bottom right rows

    #TopRight
    PlotsAdjustedM[i,8]<- (PlotsSquareM[i,8]*cos(Theta))+ (PlotsSquareM[i,9]*sin(Theta)) + NorthingCorner_0.0   # Y top right ranges
    PlotsAdjustedM[i,9]<- (PlotsSquareM[i,9]*cos(Theta))- (PlotsSquareM[i,8]*sin(Theta)) + EastingCorner_0.0  # X top right rows

    #TopLeft
    PlotsAdjustedM[i,10]<- (PlotsSquareM[i,10]*cos(Theta))+ (PlotsSquareM[i,11]*sin(Theta)) + NorthingCorner_0.0   # Y top left ranges
    PlotsAdjustedM[i,11]<- (PlotsSquareM[i,11]*cos(Theta))- (PlotsSquareM[i,10]*sin(Theta)) + EastingCorner_0.0  # X top left rows

    ##BottomLeft Adjusted Buffer
    PlotsAdjustedMBuf[i,4]<- (PlotsSquareMBuf[i,4]*cos(Theta))+ (PlotsSquareMBuf[i,5]*sin(Theta)) + NorthingCorner_0.0 # Y bottom left ranges
    PlotsAdjustedMBuf[i,5]<- (PlotsSquareMBuf[i,5]*cos(Theta))- (PlotsSquareMBuf[i,4]*sin(Theta)) + EastingCorner_0.0  # X bottom left rows

    #BottomRight Adjusted Buffer
    PlotsAdjustedMBuf[i,6]<- (PlotsSquareMBuf[i,6]*cos(Theta))+ (PlotsSquareMBuf[i,7]*sin(Theta)) + NorthingCorner_0.0# Y bottom right ranges
    PlotsAdjustedMBuf[i,7]<- (PlotsSquareMBuf[i,7]*cos(Theta))- (PlotsSquareMBuf[i,6]*sin(Theta)) + EastingCorner_0.0 # X bottom right rows

    #TopRight Adjusted Buffer
    PlotsAdjustedMBuf[i,8]<- (PlotsSquareMBuf[i,8]*cos(Theta))+ (PlotsSquareMBuf[i,9]*sin(Theta)) + NorthingCorner_0.0# Y top right ranges
    PlotsAdjustedMBuf[i,9]<- (PlotsSquareMBuf[i,9]*cos(Theta))- (PlotsSquareMBuf[i,8]*sin(Theta)) + EastingCorner_0.0  # X top right rows

    #TopLeft Adjusted Buffer
    PlotsAdjustedMBuf[i,10]<- (PlotsSquareMBuf[i,10]*cos(Theta))+ (PlotsSquareMBuf[i,11]*sin(Theta)) + NorthingCorner_0.0# Y top left ranges
    PlotsAdjustedMBuf[i,11]<- (PlotsSquareMBuf[i,11]*cos(Theta))- (PlotsSquareMBuf[i,10]*sin(Theta)) + EastingCorner_0.0  # X top left rows

    }
    }
  }
  names(PlotsAdjustedM) <- c( "Plot", "Range", "Row", "YBL", "XBL", "YBR", "XBR", "YTR", "XTR", "YTL", "XTL")
  names(PlotsAdjustedMBuf) <- c( "Plot", "Range", "Row", "YBLbuf", "XBLbuf", "YBRbuf", "XBRbuf", "YTRbuf", "XTRbuf", "YTLbuf", "XTLbuf")


  ###### PLOTTING SIZE OF PLOT DIMENSIONS ############
  if (RotatePlot==T){
  PlotY<- cbind(PlotsAdjustedM[,4], PlotsAdjustedM[,6], PlotsAdjustedM[,8], PlotsAdjustedM[,10])
  PlotX<- cbind(PlotsAdjustedM[,5], PlotsAdjustedM[,7], PlotsAdjustedM[,9], PlotsAdjustedM[,11])

  ###### PLOTTING SIZE OF PLOT DIMENSIONS ############
  PlotYBuf<- cbind(PlotsAdjustedMBuf[,4], PlotsAdjustedMBuf[,6], PlotsAdjustedMBuf[,8], PlotsAdjustedMBuf[,10])
  PlotXBuf<- cbind(PlotsAdjustedMBuf[,5], PlotsAdjustedMBuf[,7], PlotsAdjustedMBuf[,9], PlotsAdjustedMBuf[,11])

  ########## PLOTTING ##############
  pdf(paste(field,outfile,"Rotated_plots.pdf",sep="_"),
      width=30,
      height=30)
  plot(c(min(PlotX),max(PlotX)), c(min(PlotY),max(PlotY)), type="n", xlab = "UTM Easting Coordinates (m)", ylab = "UTM Northing Coordinates (m)") # Plot lines in field

  for (i in 1:nrow(PlotsAdjustedM)) {
    for (j in 1:4) {
      points(PlotX[i,j], PlotY[i,j])
    }
    polygon(PlotX[i,], PlotY[i,])
    polygon(PlotXBuf[i,], PlotYBuf[i,], col="RED")
    text( ((PlotX[i,1]+PlotX[i,3])/2), ((PlotY[i,1]+PlotY[i,3])/2), labels= PlotsSquareM[i,1], pos = NULL, col = "BLUE", cex=0.7, srt=srt_theta)
  }
  dev.off()
  }


 ###################################################################
#   ########## Make Shape Files4 ##############
# if(multirowind!=T){

  # if (file.exists(paste(output,"\\",field,".shp",sep="")) |
  #     file.exists(paste(output,"\\",field,".dbf",sep="")) |
  #     file.exists(paste(output,"\\",field,".shx",sep="")))
  #   {
  #   file.remove(paste(output,"\\",field,".shp",sep=""))
  #   file.remove(paste(output,"\\",field,".dbf",sep=""))
  #   file.remove(paste(output,"\\",field,".shx",sep=""))
  #   }


  PolygonsToMake <- vector('list', length(nPlot))
  for (i in 1: nrow(PlotsAdjustedM)) {
    PolygonsToMake[[i]] <- Polygons(list(Polygon(matrix(c(PlotsAdjustedM[i,c(5,7,9,11,5)],
                                                          PlotsAdjustedM[i,c(4,6,8,10,4)]),5,2),
                                                 hole=FALSE)),
                                    rownames(PlotsAdjustedM)[i])
  }

  if(is.null(UTMzone)){
    SpatialPolygonsToMake <- SpatialPolygons((PolygonsToMake))
    warning("Coordinate reference system not defined 'UTMzone=NULL'; This may result in difficulties loading shapefiles into other programs.")
  }
  
  if(!is.null(UTMzone)&Hemisphere=="N"){
    SpatialPolygonsToMake <- SpatialPolygons((PolygonsToMake),
                                             proj4string = CRS(
                                               as.character(paste("+proj=utm +zone=",UTMzone," +datum=NAD83 +units=m +no_defs +ellps=GRS80",sep=""))  
                                             ))
  }
  
  if(!is.null(UTMzone)&Hemisphere=="S"){
    SpatialPolygonsToMake <- SpatialPolygons((PolygonsToMake),
                                             proj4string = CRS(
                                               as.character(paste("+proj=utm +zone=",UTMzone," +south +datum=NAD83 +units=m +no_defs +ellps=GRS80",sep=""))  
                                             ))
  }

  SpatialPolygonsToMake.df <- SpatialPolygonsDataFrame(SpatialPolygonsToMake,
                                                       data.frame(id=rownames(PlotsAdjustedM),
                                                                  row.names=rownames(PlotsAdjustedM)))


  FolderpathGPSPlots<-paste(field,"_",outfile,".shp",sep="")
  writeOGR(SpatialPolygonsToMake.df,
           FolderpathGPSPlots,
           field,
           verbose = TRUE,
           overwrite_layer = T,
           driver="ESRI Shapefile")


  ########## Make Shape Files4 ##############
  # ord_infile<-infile[order(infile$Range,infile$Row)]

  # IDs<- as.character(infile$Barcode)

  PolygonsToMake <- vector('list', length(nPlot))
  for (i in 1: nrow(PlotsAdjustedM)) {
    PolygonsToMake[[i]] <- Polygons(list(Polygon(matrix(c(PlotsAdjustedMBuf[i,c(5,7,9,11,5)],
                                                          PlotsAdjustedMBuf[i,c(4,6,8,10,4)]),5,2), hole=FALSE)),
                                    rownames(PlotsAdjustedMBuf)[i])
  }

  if(is.null(UTMzone)){
    SpatialPolygonsToMake <- SpatialPolygons((PolygonsToMake))
    warning("Coordinate reference system not defined 'UTMzone=NULL'; This may result in difficulties loading shapefiles into other programs.")
  }
  
  if(!is.null(UTMzone)&Hemisphere=="N"){
    SpatialPolygonsToMake <- SpatialPolygons((PolygonsToMake),
                                             proj4string = CRS(
                                               as.character(paste("+proj=utm +zone=",UTMzone," +datum=NAD83 +units=m +no_defs +ellps=GRS80",sep=""))  
                                             ))
  }
  
  if(!is.null(UTMzone)&Hemisphere=="S"){
    SpatialPolygonsToMake <- SpatialPolygons((PolygonsToMake),
                                             proj4string = CRS(
                                               as.character(paste("+proj=utm +zone=",UTMzone," +south +datum=NAD83 +units=m +no_defs +ellps=GRS80",sep=""))  
                                             ))
  }

  SpatialPolygonsToMake.df <- SpatialPolygonsDataFrame(SpatialPolygonsToMake,
                                                       data.frame(id=rownames(PlotsAdjustedMBuf),
                                                                  row.names=rownames(PlotsAdjustedMBuf)))


  FolderpathGPSPlots<-paste(field,outfile,"buff.shp",sep="_")
  writeOGR(SpatialPolygonsToMake.df,
           FolderpathGPSPlots,
           paste(field,"Buff",sep="_"),
           verbose = TRUE,
           overwrite_layer = T,
           driver="ESRI Shapefile")

  
}
  # }
```
