## Supplemental File 1 for "R/UAStools::plotshpcreate: Create Multi-Polygon Shapefiles for Extraction of Research Plot Scale Agriculture Remote Sensing Data": SampleInfile.R

```
#' @title Infile Sample Dataset
#'
#' @description Example of infile structure for the plotshpcreat function.
#' Contains the preperation file used to design the CS17-G2FE experiment, which
#' contains 594 two row plots and 1188 observations.
#'
#' @name SampleInfile
#' @docType data
#' @usage SampleInfile
#' @format A dataframe containg the preperation file used to design the CS17-G2FE experiment, which
#' contains 594 two row plots and 1188 observations.
#'
#' We recommend have repeated Barcodes and Plot numbers if there are multirow plots (mrowplot>1) as
#' the plotshpcreatre function accounts for this redundance within the function.
#'  [,1]  Loc         An abbreviation to identify the trial location and year (character)
#'  [,2]  Test        An abbreviation to define the trial name  (character)
#'  [,3]  Plot        The number of each plot (numeric)
#'  [,4]  Barcode     A unique identifier for each plot (character)
#'  [,5]  Stock       The seed source which is planted in each plot (character)
#'  [,6]  Pedigree    Information defining the lineage of the seed source (character)
#'  [,7]  Entry       An additional unique identifer commonly used to track consistent pedigrees from different seed stocks (numeric)
#'  [,8]  Rep         The replicate number of each plot (numeric)
#'  [,9]  Range       The range [also called row] number of each plot in the experimental grid (numeric)
#'  [,10] Row         The row [also called column] number of each plot in the experimental grid (numeric)
#' @note
#' Only Range, Row, Plot, and Barcode variables are necessary for the the proper execution of plotshpcreate,
#' but additional columns may be included in the dataframe.
#' @keywords datasets infile sample
NULL
```
