## Supplemental File 1 for "R/UAStools::plotshpcreate: Create Multi-Polygon Shapefiles for Extraction of Research Plot Scale Agriculture Remote Sensing Data": README.md

# [R/UAStools](https://github.com/andersst91/UAStools/wiki) 

### Authors:
Steven L. Anderson II  
Seth C. Murray

### Description:
UAStools is a package developed to provide useful tools for working with remote sensing data set, specifically in reference to field-based, agriculture research plots. Currently, the [plotshpcreate.R](https://github.com/andersst91/UAStools/wiki/plotshpcreate.R) function is available to construct a multi-polygon shapefile (.shp) of a reasearch trial, with individual polygons defining specific research plot.


# Installation  
Make sure you have the latest version of [R](https://cran.r-project.org/). 

[R/UAStools](https://github.com/andersst91/UAStools/wiki) is availible on [GitHub](https://github.com/andersst91/UAStools).

You then need to install the [devtools](https://github.com/r-lib/devtools) package, plus a set of package dependencies: [sp](https://cran.r-project.org/web/packages/sp/index.html) and [rgdal](https://cran.r-project.org/web/packages/rgdal/index.html). (Additional, secondary dependencies will also be installed.)

To install [R/UAStools](https://github.com/andersst91/UAStools/wiki) execute the following commands in R:  

    install.packages("devtools") 
    library("devtools")
    devtools::install_github("andersst91/UAStools")
    library("UAStools")

# Availible Functions
## [plotshpcreate.R](https://github.com/andersst91/UAStools/wiki/plotshpcreate.R)  
Construct a multi-polygon shapefile (.shp) of a reasearch trial, with individual polygons defining specific research field plots.  
Visit the wiki page for more details: https://github.com/andersst91/UAStools/wiki/plotshpcreate.R  

### Note: Make sure to setwd() before executing [plotshpcreate.R](https://github.com/andersst91/UAStools/wiki/plotshpcreate.R)!!!

    ?plotshpcreate
    ?SampleInfile
    
### Citation

To cite R/UAStools in publications, use:  

Anderson, S.L., S. Murray, L. Malambo, C. Ratcliff, S. Popescu, D. Cope, et al. 2019. Prediction of Maize Grain Yield Before Maturity Using Improved Temporal Height Estimates of Unmanned Aerial Systems. The Plant Phenome Journal. doi:10.2135/tppj2019.02.0004

### Liscense
[Licensed](LICENSE) under [GPL-2](https://www.r-project.org/Licenses/GPL-2).
